# Insulin promoted mobilization of GLUT4 from a perinuclear storage site requires RAB10

**DOI:** 10.1101/2020.04.14.040683

**Authors:** Alexandria Brumfield, Natasha Chaudhary, Dorothee Molle, Jennifer Wen, Johannes Graumann, Timothy E. McGraw

## Abstract

Insulin controls glucose uptake into muscle and fat cells by inducing a net redistribution of GLUT4 from intracellular storage to the plasma membrane (PM). The TBC1D4-RAB10 signaling module is required for insulin-stimulated GLUT4 translocation to the PM, although where it intersects GLUT4 traffic was unknown. Here we demonstrate that TBC1D4-RAB10 functions to control GLUT4 mobilization from a Trans Golgi Network (TGN) storage compartment, establishing that insulin, in addition to regulating the PM proximal effects of GLUT4-containing vesicles docking to and fusion with the PM, also directly regulates the behavior of GLUT4 deeper within the cell. We also show that GLUT4 is retained in an element/domain of the TGN from which newly synthesized lysosomal proteins are targeted to the late endosomes and the ATP7A copper transporter is translocated to the PM by elevated copper. Insulin does not mobilize ATP7A nor does copper mobilize GLUT4. Consequently, GLUT4 intracellular sequestration and mobilization by insulin is achieved, in part, through utilizing a region of the TGN devoted to specialized cargo transport in general rather than being specific for GLUT4. Our results define GLUT4-containing region of the TGN as a sorting and storage site from which different cargo are mobilized by distinct signals.

## INTRODUCTION

Regulation of glucose uptake by fat and muscle cells, essential for the maintenance of whole-body glucose homeostasis, is determined by the levels of glucose transporter 4 (GLUT4) in the plasma membranes (PM) of these cells (1). GLUT4 cycles between intracellular compartments and the PM, with the distribution determined by the rates of exocytosis and endocytosis (2–5). The main effect of insulin is to stimulate GLUT4 exocytosis to increase the amount of PM GLUT4, thereby promoting increased glucose uptake (1).

In the basal state (unstimulated cells) the majority of GLUT4 resides intracellularly in perinuclear compartments that are in part Trans Golgi Network (TGN) in nature (6–8), and in specialized vesicles (referred to as insulin-responsive vesicles, IRVs) dispersed throughout the cytosol (9–11) whose delivery to the PM is regulated by insulin (2). GLUT4 in the PM cycles back to the TGN via the endosomal pathway (2, 12, 13). Targeting GLUT4 from endosomes to the TGN has an important role in basal intracellular GLUT4 retention. Mutations in GLUT4 that disrupt its traffic from endosomes to the TGN are poorly retained in basal conditions and are not properly translocated to the PM upon insulin stimulation (6, 14–16). These results identify the TGN as the site for formation of IRVs. The TGN is a main sorting compartment along the biosynthetic and endocytic pathways. Cargoes to be targeted to distinct destinations are sorted and packaged into the correct transport vesicles in the TGN. The relationship between the TGN containing GLUT4 and the TGN involved in the traffic of other cargoes is not known (2, 6, 8, 12).

Insulin signaling triggers multiple discrete molecular events that mediate efficient recruitment, docking, and fusion of IRVs with the PM (17–19). These events lead to a decrease in the size of the intracellular GLUT4 pool concomitant with an increase of GLUT4 in the PM. As GLUT4 in the PM is in equilibrium with intracellular GLUT4, endocytosis of GLUT4 dynamically removes GLUT4 from the PM. Thus, maintenance of the insulin-stimulated dynamic increase in GLUT4 in the PM requires the continual ferrying of GLUT4-containing IRVs to the PM. Insulin signaling can add to the IRV pool by increasing the rate of GLUT4 mobilization from the TGN in nascent IRVs. Despite the biological importance, insulin regulation of GLUT4 trafficking at the perinuclear region has not been thoroughly interrogated.

A key aspect of insulin regulation of GLUT4 trafficking is inhibition of the GTPase-activating protein (GAP) TBC1D4/AS160 (20, 21), allowing for activation of its target RAB, RAB10 (22). In 3T3-L1 adipocytes and primary adipocytes, knockdown of TBC1D4 releases the inhibition of GLUT4 exocytosis in the basal state by 50% (21, 23), and depletion of RAB10 in cultured and primary adipocytes specifically blunts insulin-stimulated GLUT4 translocation by 50% (22, 24, 25). These data demonstrate insulin-stimulated GLUT4 exocytosis is regulated by both TBC1D4-RAB10-dependent and independent mechanisms. Overexpressed RAB10 has been shown to reside on IRVs in adipocytes (13), and total internal reflection fluorescence (TIRF) microscopy studies have demonstrated RAB10 functions at a step prior to IRV fusion with the PM (24). Thus, it is commonly thought RAB10 regulates IRV recruitment and/or docking with the PM. In other cell types RAB10 is required for regulated trafficking processes that involve vesicle delivery to the PM (26–30), and RAB10 has been shown to localize to both perinuclear and vesicular compartments (29, 30).

We have previously identified SEC16A as a novel RAB10-interacting protein required for insulin-stimulated GLUT4 translocation (31). Knockdown of SEC16A in 3T3-L1 adipocytes specifically blunts insulin-stimulated GLUT4 translocation by 50%, with no additivity of double knockdown of RAB10 and SEC16A (31). Interestingly, a pool of SEC16A localizes to structures in the perinuclear region that encircle GLUT4-containing perinuclear membranes (31). SEC16A is known to localize to endoplasmic reticulum exit sites (ERES) and act as a scaffold for organization of COPII components required for budding of COPII vesicles (32, 33). In adipocytes, SEC16A’s role in GLUT4 trafficking is independent of its role in ERES function since knockdown of other components of the ER exit site machinery, which blunt secretion, are without effect on GLUT4 translocation (31). SEC16A’s perinuclear localization, and lack of SEC16A localization to IRVs (31), suggests RAB10 might function at the perinuclear region to regulate GLUT4 trafficking. In this study we use an novel proteomic approach to demonstrate that GLUT4 resides in a region of TGN where specialized cargoes are sorted and mobilized by specific stimuli, and using a novel live-cell imaging assay we demonstrate insulin promotes the mobilization of GLUT4 from the TGN through RAB10 activity.

## RESULTS

### GLUT4 is retained in a region of the TGN from which specialized cargoes are sorted and mobilized

The first approach we took to gain insight to GLUT4 trafficking at the perinuclear region was to identify proteins that reside with GLUT4 in the perinuclear compartment of 3T3-L1 adipocytes. It has previously been shown that mutation of the phenylalanine of the GLUT4 amino terminal F^5^QQI motif (amino acid positions 5 through 8) to a tyrosine (Y^5^QQI) redistributes GLUT4 to the TGN perinuclear compartment from cytosolic puncta (14). HA-GLUT4-GFP is a reporter extensively used in studies of GLUT4 traffic (34). HA-GLUT4-GFP with a F^5^QQI to Y^5^QQI mutation (F^5^Y-HA-GLUT4-GFP) displays enhanced intracellular retention in the TGN, as demonstrated by increased colocalization with TGN markers Syntaxin6 (STX6) and TGN46 (Fig. 1A). F^5^Y-GLUT4 continually cycles to and from the PM, and thus is dynamically concentrated in the TGN (14). GLUT4 in which an alanine is substituted for phenylalanine in the F^5^QQI motif (F^5^A-GLUT4) is not as efficiently targeted to the TGN as compared to WT GLUT4 (14–16). Consequently, F^5^A-HA-GLUT4-GFP was predominantly localized in vesicular elements throughout the cytoplasm and was not well concentrated around the nucleus (Fig. 1A). Given F^5^Y-GLUT4 is enriched in the TGN as compared to both WT GLUT4 and F^5^A-GLUT4 (14), we reasoned proteins colocalized with GLUT4 in the TGN (the same TGN membrane as GLUT4) would be enriched in a detergent-free immunoabsorption of F^5^Y-GLUT4 because membrane integrity is preserved in this protocol. Membrane compartments containing HA-GLUT4-GFP were isolated by detergent-free immunoabsorption with anti-GFP-antibody from mechanically-disrupted unstimulated 3T3-L1 adipocytes stably expressing HA-GLUT4-GFP, F^5^Y-HA-GLUT4-GFP or F^5^A-GLUT4-GFP. We used stable isotope labelling with amino acids in culture (SILAC) to quantitatively compare by mass spectrometry proteins co-immunoabsorbed with these different GLUT4 constructs (35, 36). Pair-wise comparisons of WT versus F^5^Y and F^5^A versus F^5^Y were performed in duplicate inverting which cells were grown in the heavy amino acid medium, generating 4 different data sets of proteins immunoabsorbed with F^5^Y compared to WT (2 sets) and compared to F^5^A (2 sets).

**Figure 1.**
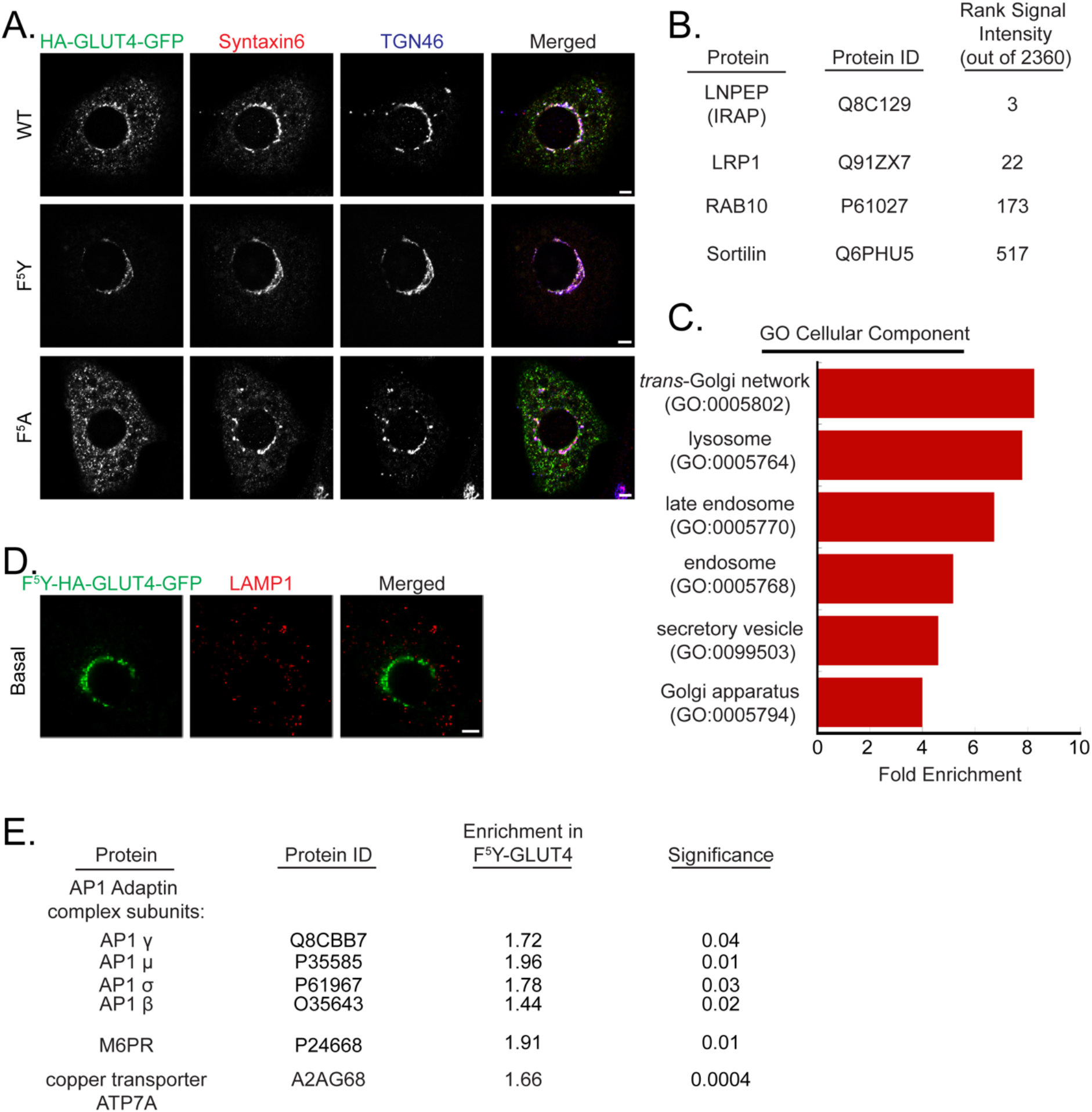
Proteomic analysis of GLUT4-containing perinuclear compartments. **A.** Representative Airyscan confocal single plane images of cells expressing wildtype (WT), F^5^Y-, or F^5^A-HA-GLUT4-GFP and labeled for Syntaxin6 and TGN46 by IF. **B.** Proteins identified in immunoabsorption experiments that are known to colocalize with GLUT4, rank based on summed signal intensity from 4 immunoabsoprtion experiments. **C.** Panther Gene Ontology (GO) cellular component analysis for localization of proteins increased in F^5^Y-GLUT4 compartments immunoadsorption. **D.** Representative Airyscan confocal single plane images of cells expressing F^5^Y-HA-GLUT4-GFP mutant and labeled for LAMP1 by IF. **E.** Fold increase of AP1 adaptin complex subunits, mannose 6-phosphate receptor (MPR), and copper transporter ATP7A in F^5^Y-GLUT4 compartments immunoadsorption. Bars, 5 μm.

There were 2360 proteins in the merged data from the 4 sets of data. The experimental premise that mechanical disruption preserves, at least partially, the integrity of membrane compartments/domains was validated by the fact that relative abundance (summed signal intensity) of proteins previously identified to colocalize but not directly interact with GLUT4, including: LNPEP (or IRAP) (37), LRP1 (9), RAB10 (13, 22) and Sortilin (38), were in the top 20% of proteins ranked based on signal intensity (Fig. 1B). Of note, LNPEP, which is known to traffic via the same pathway as GLUT4 (37) and is therefore expected to be efficiently co-immunoabsorbed with GLUT4, was the 3^rd^ most abundant protein in the immunoabsorption based on signal intensity.

The aim of this study was to identify proteins enriched in the F^5^Y-GLUT4 immunoabsorption; therefore, we focused our analyses on the set of 508 proteins that in the pooled data set were increased in the F^5^Y-GLUT4 immunoabsorption by greater than 1.3 fold by SILAC-ratio. Gene ontology cellular component analyses (39, 40) revealed a significant enrichment for proteins annotated to be localized to the TGN and TGN transport vesicles, including STX6 and TGN46 (Fig. 1C). In addition, there was enrichment of Golgi, endosome, exocytic vesicles and the ER-to-Golgi intermediate compartment proteins (Fig. 1C), consistent with GLUT4 being dynamically distributed among a number of intracellular compartments (2).

Unexpectedly, there was also a significant enrichment of late endosome and lysosome proteins (Fig. 1C). This enrichment was not because F^5^Y-GLUT4 is localized to late endosomes/lysosomes as there is no significant colocalization between F^5^Y-GLUT4 and LAMP1 (Fig. 1D). The majority of newly synthesized lysosomal proteins are delivered to the lysosomes by a pathway involving targeting from the TGN to the late endosomes (41). Soluble lysosome proteins, which are modified by mannose 6-phosphate in the ER, are diverted from delivery to the PM at the level of the TGN via a mechanism requiring the mannose 6-phosphate receptor (MPR) and the AP1 clathrin adaptin complex (41). Thus, an explanation for the enrichment of F^5^Y-GLUT4 with lysosomal proteins is that the GLUT4-containing perinuclear concentration is a specialized sub compartment of the TGN where lysosomal proteins are diverted from delivery to the PM by sorting to specialized transport vesicles. In support of that hypothesis, the MPR and the 4 subunits of the AP1 complex (AP1 μ,σ,ß,γ), components of the machinery that targets lysosomal proteins to the late endosomes, were significantly enriched in the F^5^Y-GLUT4 immunoabsorption (Fig. 1E). Based on these data we propose that the region of the TGN enriched for F^5^Y-GLUT4 is involved in the sorting of cargoes that exit the TGN via specialized vesicles, diverting cargo from non-specialized vesicles that mediate constitutive traffic to the PM.

The immunoabsorption data identified the Menkes copper transporter, ATP7A, as enriched in the F^5^Y-GLUT4-containing perinuclear compartments (Fig 1E). Previous studies have also identified that ATP7A co-immunoabsorbs with GLUT4 (11). ATP7A, which is expressed in a broad variety of cell types, has a role in protecting cells against copper overload. At physiological copper levels ATP7A primarily localizes to the TGN, but with an increased copper load ATP7A translocates to the PM, where it pumps copper from cells (42). In low copper conditions, achieved by treatment with copper chelator bathocuproinedisulfonic acid disodium salt (BCS), ATP7A was predominantly colocalized with GLUT4 in the TGN of 3T3-L1 adipocytes, validating the mass spectrometry data (Fig. 2A & B). Challenging cells with elevated copper resulted in a decrease in the intensity of ATP7A in STX6-postive TGN and an increase in ATP7A labeling in cytosolic vesicles (Fig. 2A). Copper mobilization of ATP7A was reflected by a significant decrease in ATP7A overlap with STX6 (Fig. 2C). Insulin stimulation in adipocytes results in translocation of GLUT4 to the PM, as measured by ratiometric analyses of the HA-GLUT4-GFP reporter (Fig. 2D). Insulin stimulation did not affect ATP7A co-localization with STX6 (Fig. 2A & C), nor did elevated copper promote GLUT4 translocation to the PM (Fig. 2A & D). Thus, despite the high degree of colocalization of ATP7A and GLUT4, their mobilizations from the TGN are linked to distinct stimuli. These data support the hypothesis that the GLUT4-containing TGN compartment is a retention and sorting hub where various stimuli mobilize specific cargo.

**Figure 2.**
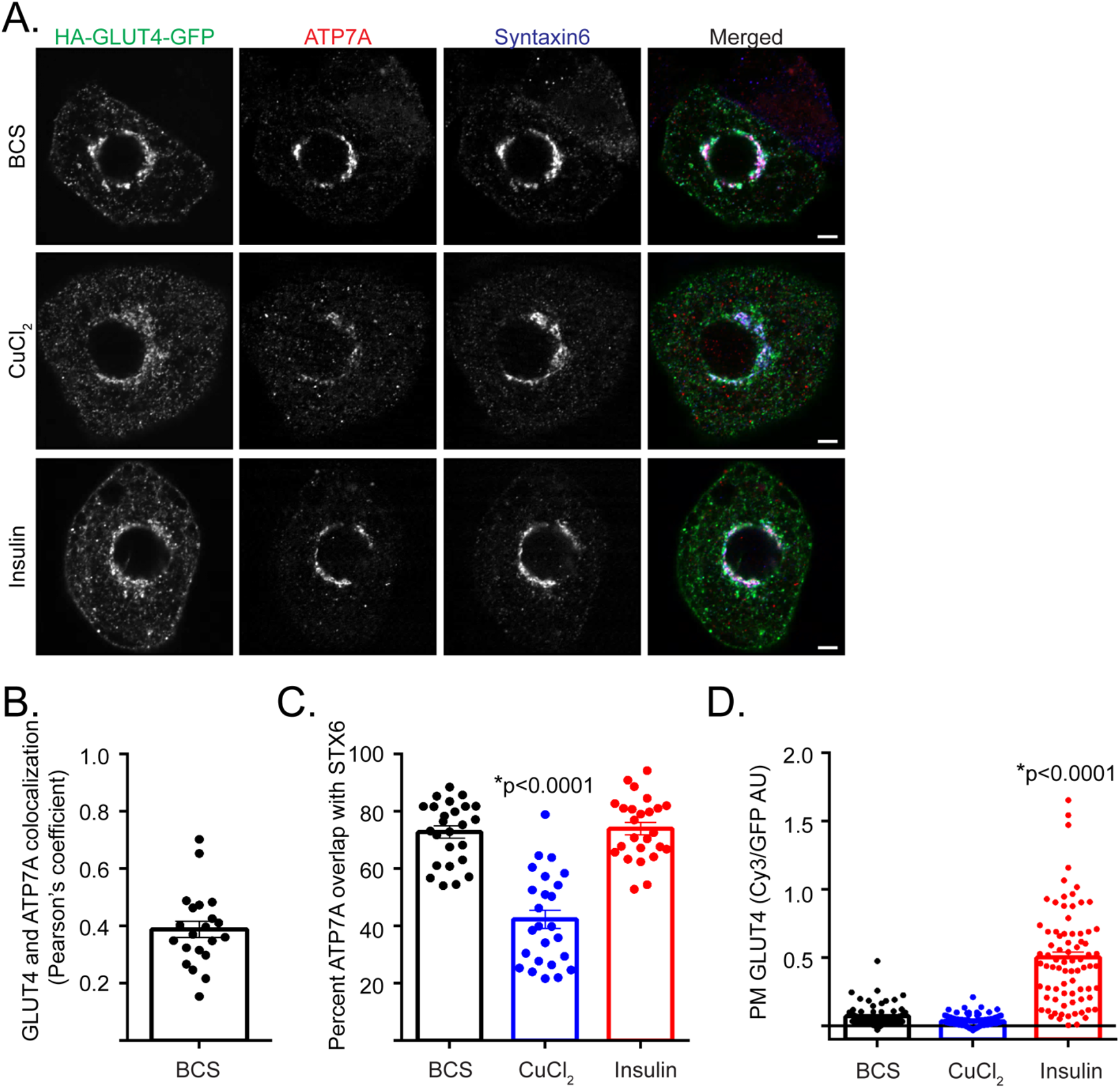
Copper, but not insulin, stimulation results in mobilization of the copper transporter ATP7A from GLUT4-containing perinuclear compartments. **A.** Representative Airyscan confocal single plane images of cells expressing HA-GLUT4-GFP and labeled for native copper transporter ATP7A and Syntaxin6 by IF. Cells treated with 200μM BCS, followed by treatment with 200μM copper or 1nM insulin as described in materials and methods. Bars, 5 μm. **B.** Pearson’s correlation coefficient (r) for colocalization between GLUT4 and ATP7A in 3T3-L1 adipocytes under BCS condition. Individual cells ± SEM from N = 3 assays. **C.** Quantification of percent overlap of ATP7A with Syntaxin6 under BCS, copper, and insulin-stimulated conditions in 3T3-L1 adipocytes. Individual cells ± SEM from N = 3 assays. **D.** Representative experiment of quantification of PM to total HA-GLUT4-GFP in cells under BCS, copper, and insulin-stimulated conditions, as described in materials and methods. Individual cells ± SEM. AU, arbitrary units. *, p<0.05 compared to BCS condition, two-tailed unpaired t-test, nonnormalized raw data.

### Insulin increases the rate of GLUT4 mobilization from the perinuclear region

We next sought to demonstrate that insulin stimulation promotes the mobilization of GLUT4 from the perinuclear region of 3T3-L1 adipocytes, similar to copper stimulation promoting the mobilization of ATP7A. With insulin stimulation it was visually apparent that the GLUT4-containing IRV pool was decreased in size concomitant with an increase in GLUT4 in the PM (Fig. 3A). However, in static images an effect of insulin on GLUT4 in the TGN was not apparent. Visualizing the mobilization of GLUT4 from the perinuclear compartment in live-cell imaging would prove very useful in determining if insulin regulates GLUT4 trafficking at the perinuclear region, yet has been confounded by the difficulty of distinguishing GLUT4-containing vesicles that have budded from the perinuclear compartments from those that have been endocytosed at the plasma membrane. To overcome this limitation we tagged GLUT4 with an irreversible green-to-red photoconvertible protein mEos3.2 (43) (HA-GLUT4-mEos3.2) and visualized the mobilization of HA-GLUT4-mEos3.2 that has been acutely photoconverted from green to red in a region of the perinuclear compartment, and thus could be distinguished from the remainder of HA-GLUT4-mEos3.2 in the cell (Fig. 3B). After photoconversion, the decrease over time in red HA-GLUT4-mEos3.2 intensity in the photoconverted region represents GLUT4 that has been mobilized from the perinuclear region, and the return over time of the green HA-GLUT4-mEos3.2 intensity in the photoconverted region represents GLUT4 that has been mobilized to the perinuclear region (Fig. 3B). Importantly, the trafficking of HA-GLUT4-mEos3.2 was similar to the well characterized HA-GLUT4-GFP reporter (Fig. 3C). Furthermore, in fixed cells successive image acquisition did not result in a decrease in the red HA-GLUT4-mEos3.2 intensity in the photoconverted region (Fig. 3D). These data argue that in live-cell imaging, any decrease in red HA-GLUT4-mEso3.2 intensity observed is not a result of photobleaching with successive image acquisition.

**Figure 3.**
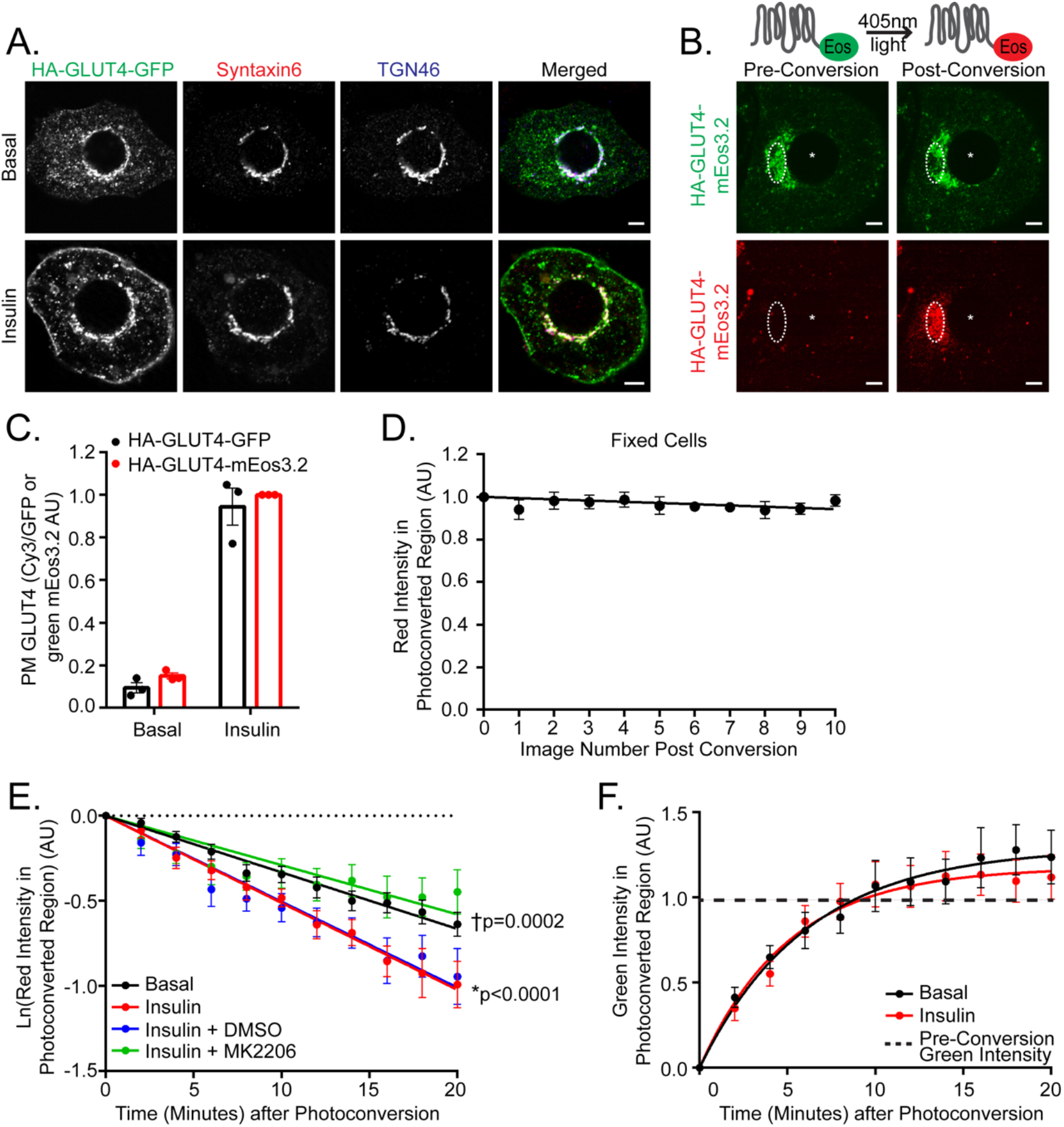
Insulin promotes mobilization of HA-GLUT4-mEos3.2 from the perinuclear region downstream of AKT. **A.** Representative Airyscan confocal single plane images of basal and insulin-stimulated cells expressing HA-GLUT4-GFP and labeled for Syntaxin6 and TGN46 by IF. **B.** Representative Airyscan confocal single plane images of cells expressing HA-GLUT4-mEos3.2. Green HA-GLUT4-mEos3.2 photoconverted to red HA-GLUT4-mEos3.2 in the perinuclear region (indicated by white, dashed circle) as described in material and methods. *, indicates nucleus. **C.** Quantification of PM to total HA-GLUT4-GFP or HA-GLUT4-mEos3.2 as described in materials and methods. Serum starved cells stimulated with 10nM insulin. Values normalized to HA-GLUT4-mEos3.2 expressing, insulin condition. N=3 assays ± SEM. **D.** Quantification of average red HA-GLUT4-mEos3.2 intensity in the photoconverted perinuclear region of fixed cells for 10 successive images. Values normalized to image 0. Mean normalized values ± SEM, N=2 assays, 6-7 cells per assay. **E.** Quantification of average red HA-GLUT4-mEos3.2 intensity in the photoconverted perinuclear region of live cells. Prior to photoconversion serum starved cells stimulated with 10nM insulin, 1μM AKT inhibitor MK2206, or equivalent volume of DMSO, where indicated, as described in materials and methods. Values normalized to value at time 0. Mean normalized values ± SEM, N=5-6 assays, 4-7 cells per assay. *, p<0.05 comparing basal and insulin-stimulated slopes. †, p<0.05 comparing insulin + DMSO and insulin + MK2206-stimulated slopes. **F.** Quantification of average green HA-GLUT4-mEos3.2 intensity in the photoconverted perinuclear region of live cells. Prior to photoconversion serum starved cells stimulated with 10nM insulin where indicated. Values normalized to value at time 0. Mean normalized values ± SEM, N=5-6 assays, 4-7 cells per assay. AU, arbitrary units. Bars, 5 μm.

We first determined if insulin regulates the mobilization of GLUT4 from the perinucelar compartment. Under basal conditions red HA-GLUT4-mEos3.2 was mobilized from the photoconverted region with a rate k=0.033 min^1^ (Fig. 3E). Under insulin-stimulated conditions red HA-GLUT4-mEos3.2 was mobilized from the photoconverted region with a rate k=0.051 min^1^ (Fig. 3E), a 1.53 fold increase compared to basal conditions. These data are the first direct evidence demonstrating that insulin signaling accelerates mobilization of GLUT4 from the perinuclear region. Insulin-stimulated GLUT4 translocation in adipocytes and muscle requires activation of AKT (1). Insulin regulation of GLUT4 mobilization from the perinuclear region is downstream of AKT in 3T3-L1 adipocytes, and as compared to insulin in the presence of DMSO (vehicle), insulin in the presence of AKT inhibitor MK2206 (44) could not promote the mobilization of HA-GLUT4-mEos3.2 from the perinuclear region (Fig. 3E).

We next determined if insulin regulates the traffic of GLUT4 to the perinuclear region. In both basal (no stimulation) and insulin-stimulated conditions the green HA-GLUT4-mEos3.2 intensity in the photoconverted region returned to the pre-photoconversion intensities with half-times of approximately 5 minutes (Fig. 3F). Thus, GLUT4 return to the TGN is not regulated by insulin. This result coupled with our finding that GLUT4 constitutively traffics from the TGN (Fig. 3E), demonstrate that GLUT4 is dynamically concentrated in the peri-nuclear region.

### RAB10 colocalizes with SEC16A and GLUT4 at the perinuclear region

To investigate if RAB10 contributes to insulin-stimulated mobilization of GLUT4 from the perinuclear region, we first determined the localization of RAB10 in 3T3-L1 adipocytes by expressing RAB10 tagged with blue fluorescent protein (BFP-RAB10) (Fig. 4A and B). A pool of RAB10 localized to the perinuclear region under basal and insulin-stimulated conditions, suggesting its perinuclear localization is independent of its GDP/GTP state (Fig. 4A and B). As demonstrated previously, perinuclear SEC16A-labeled structures encircled HA-GLUT4-GFP-containing perinuclear TGN membranes under basal and insulin-stimulated conditions (31) (Fig. 4A and B). Perinuclear BFP-RAB10 colocalized with both perinuclear SEC16A and HA-GLUT4-GFP under basal and insulin-stimulated conditions, as demonstrated by linescan analyses (Fig. 4A and B). In the context of the known functional role of RAB10 and SEC16A in GLUT4 trafficking, these data raise the possibility that RAB10 and SEC16A function at the perinuclear region to regulate GLUT4 trafficking.

**Figure 4.**
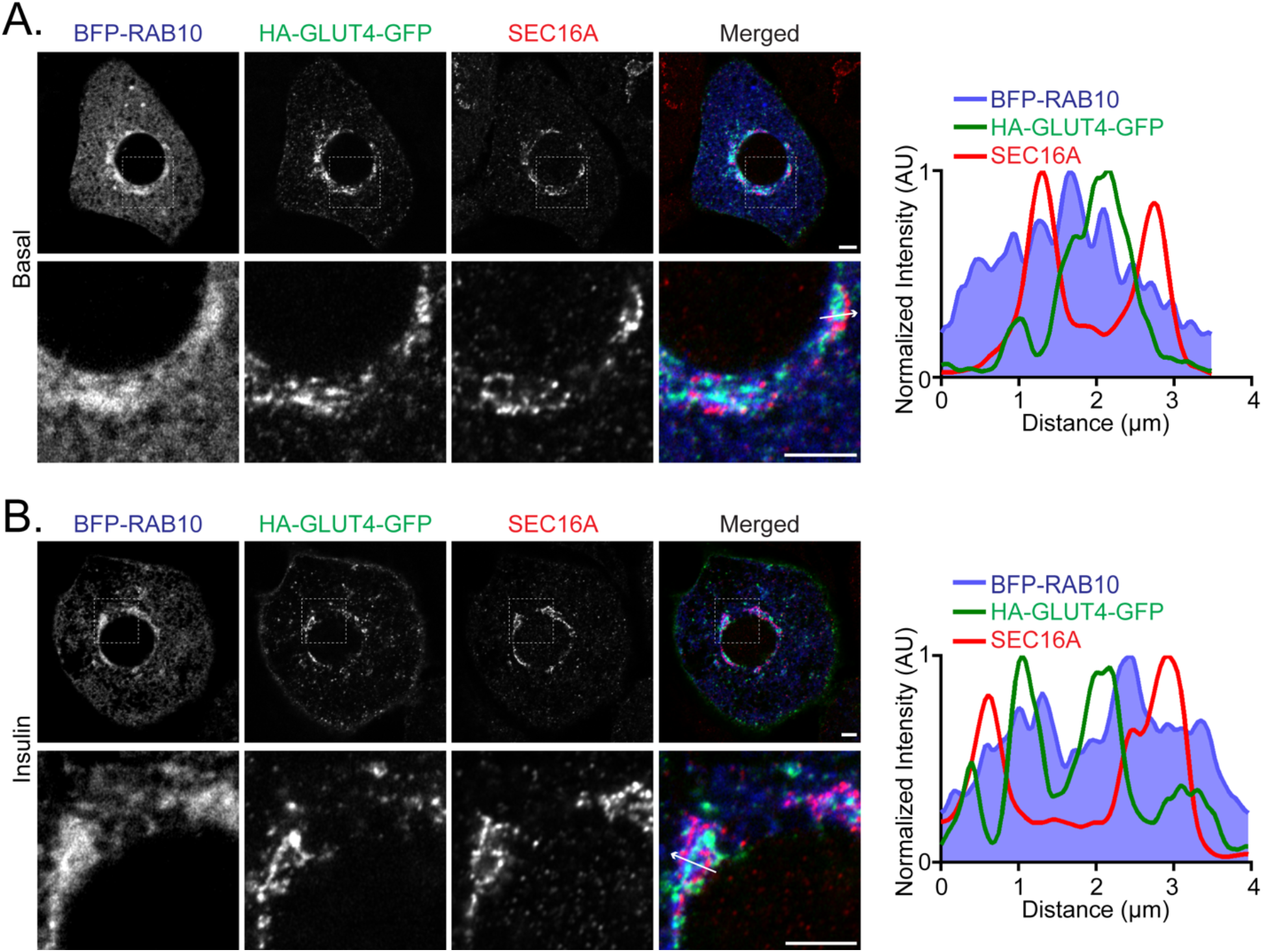
RAB10 colocalizes with HA-GLUT4-GFP and SEC16A at the perinuclear region. **A and B.** Representative Airyscan confocal single plane images of (A) basal and (B) insulin-stimulated cells expressing BFP-RAB10 and HA-GLUT4-GFP, and labeled for endogenous SEC16A by IF. Serum starved cells stimulated with 1nM insulin. Inset (white, dashed boxed region) displayed below. Linescan plot is BFP-RAB10, HA-GLUT4-GFP, and SEC16A fluorescence intensity along a line (indicated by white arrow). Values normalized to each individual fluorescence maxima. Bars, 5 μm.

### The organization of RAB10-labeled, SEC16A-labeled, and GLUT4-containing perinuclear membranes is not random

The perinuclear region is compact in nature and contains a number of different membrane compartments (i.e. Golgi, ER-to-Golgi intermediate compartments (ERGIC), and ER). Thus, we sought to determine if the spatial organization of RAB10-labeled, SEC16A-labeled, and GLUT4-containing perinuclear membranes is simply due to this compact nature, or if their spatial organization is not random and is important to the function of RAB10 and SEC16A in GLUT4 trafficking. To gain insight into this question we treated cells with nocodazole and determined if the RAB10-SEC16A-GLUT4 spatial organization is retained (Fig. 5A and B). The organization of the Golgi as a ribbon-like organelle and its perinuclear localization is highly dependent on an intact microtubule cytoskeleton (45, 46). Nocodazole-induced disruption of microtubule polymerization leads to fragmentation and dispersion of the Golgi throughout the cytosol (45, 46). When the Golgi fragments, Golgi ministacks are formed that retain the structural polarity of the *cis-, medial-,* and *trans*-Golgi. The Golgi ministacks are recapitulated at peripheral endoplasmic reticulum exit sites (ERES) to re-establish ER to Golgi secretion (45). With nocodazole treatment, we observed that the spatial organization of RAB10, SEC16A, and GLUT4 described above was retained under basal and insulin-stimulated conditions (Fig. 5A and B). By performing a radial line scan analysis centered on HA-GLUT4-GFP, we demonstrated SEC16A-labeled membranes remained adjacent to HA-GLUT4-GFP-containing membranes, and RAB10 remained localized with both SEC16A and GLUT4 (Fig. 5C and D). Furthermore, the average distance between peaks of HA-GLUT4-GFP fluorescence and SEC16A fluorescence was approximately 800nm in both the presence and absence of nocodazole (Fig. 5E). These data suggest the RAB10-SEC16A-GLUT4 perinuclear organization is not random and could be important for RAB10-SEC16A function in GLUT4 trafficking.

**Figure 5.**
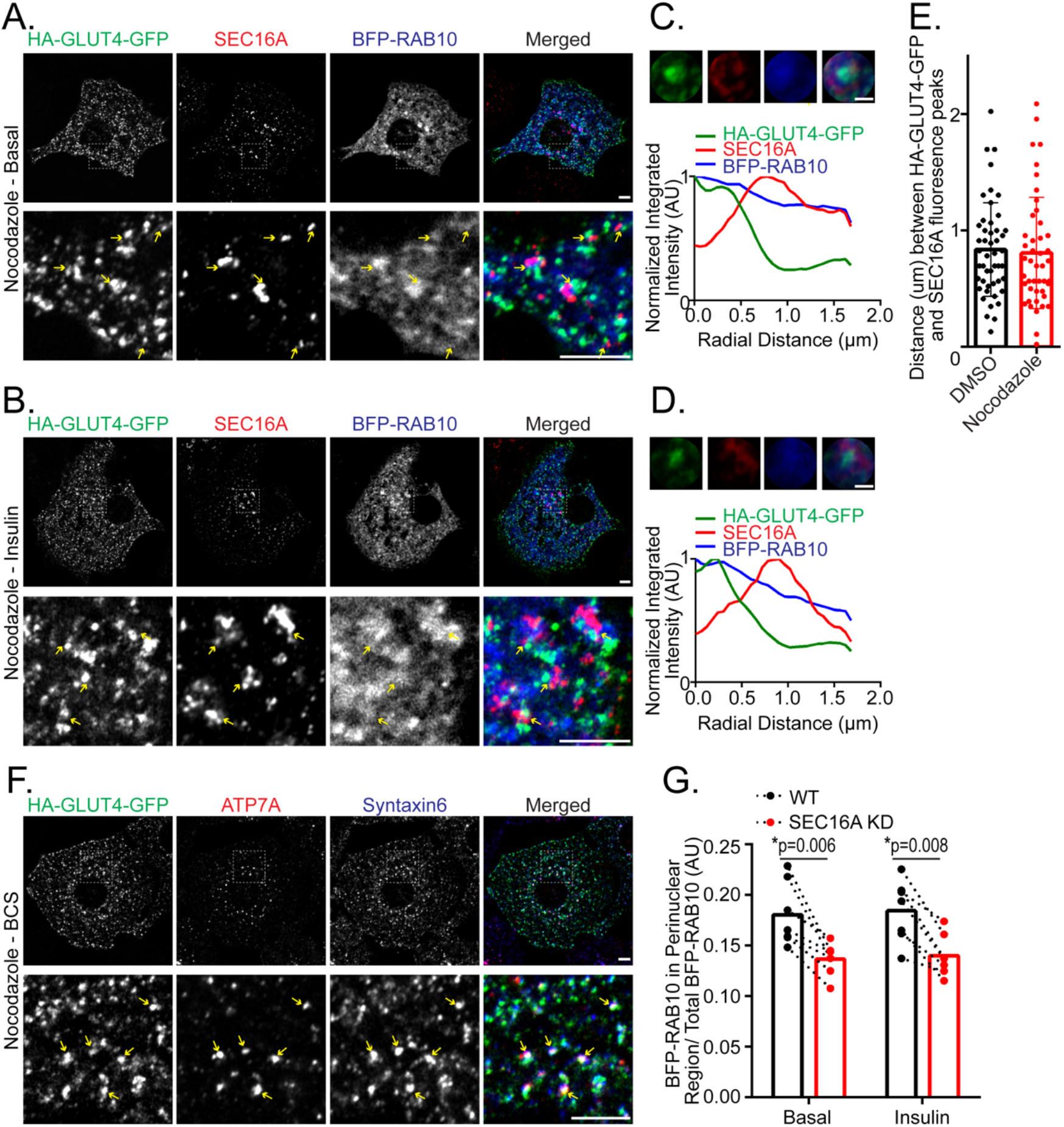
The organization of perinuclear RAB10 and SEC16A with GLUT4 has implications for their function in GLUT4 trafficking. **A and B.** Representative Airyscan confocal single plane images of cells treated with 3μM nocodazole. Cells expressing HA-GLUT4-GFP and BFP-RAB10 and stained for endogenous SEC16A by IF. Cells under (A) basal and (B) 1nM insulin-stimulated conditions. Inset (white, dashed boxed region) displayed below. Yellow arrows indicate the same position in each image. Bars, 5 μm. **C and D.** Images of the average HA-GLUT4-GFP, SEC16A, and BFP-RAB10 fluorescence intensity from 5 individual fragments, centered of HA-GLUT4-GFP, resulting from nocodazole treatment from the cells in A and B respectively. Radial linescan plot of images displayed below. Values normalized to each individual fluorescence maxima. Bars, 1 μm. **E.** Quantification of the distance (μm) between HA-GLUT4-GFP and SEC16A fluorescence peaks in basal cells in the presence and absence of nocodazole treatment. Values are distances between peaks ± SEM. Distance measured for 3 separate sets of peaks per cell. N=2 assays, 7-8 cells per assay. **F.** Representative Airyscan confocal single plane images of cells treated with 3μM nocodazole in the presence of 200μM BCS. Cells expressing HA-GLUT4-GFP and stained for endogenous ATP7A and Syntaxin6 by IF. Inset (white, dashed boxed region) displayed below. Yellow arrows indicate the same position in each image. Bars, 5 μm. **G.** Quantification of the fraction of BFP-RAB10 in the perinuclear region of basal and 1nM insulin-stimulated cells ± addition of siRNA targeting SEC16A. N=7 assays ± SEM. Dashed line connects data from individual assays. *, p<0.05, two-tailed unpaired t-test, nonnormalized raw data.

Given the organization of perinuclear RAB10-SEC16A-GLUT4 is retained with nocodazole treatment, we reasoned the colocalization of cooper transporter ATP7A with GLUT4 at the TGN (Fig. 2A and B) should be retained in fragments formed with nocodazole treatment. Indeed, we observed with nocodazole treatment that ATP7A colocalized with GLUT4 in a subset of GLUT4-containing fragments that contain Syntaxin6 (Fig. 5F).

### Perinuclear SEC16A is important for proper localization of RAB10 at the perinuclear region

Given SEC16A is known to act as a scaffold for organization of COPII components at ERES (32, 33), we wondered whether perinuclear SEC16A analogously acts as a scaffold for organization of RAB10 at perinuclear membranes. We found 18% of total BFP-RAB10 localized to the perinuclear region under basal and insulin-stimulated conditions (Fig. 5G). Depletion of SEC16A resulted in a 30% decrease in BFP-RAB10 in the perinuclear region under basal and insulin-stimulated conditions (Fig. 5G), demonstrating the presence of the perinuclear pool of SEC16A is important for localizing RAB10 at perinuclear membranes. These data argue SEC16A-dependent localization of RAB10 at perinuclear membranes is independent of its GTP/GDP status, and if properly localized at the perinuclear region RAB10 bound to GTP can carry out its function in insulin-stimulated GLUT4 trafficking.

### The RAB10-AS160 module regulates GLUT4 mobilization from the perinuclear region

We next determined if RAB10 and its GAP TBC1D4 regulate the rate of GLUT4 mobilization from the perinuclear region. Depletion of TBC1D4 results in constitutive activation of RAB10 (22), and knockdown of TBC1D4 in the absence of insulin stimulation led to acceleration of the mobilization of red HA-GLUT4-mEos3.2 from the perinuclear region near to the insulin-stimulated rate (Fig. 6A). The effect of TBC1D4 depletion on mobilization of HA-GLUT4-mEos3.2 was rescued by expression of shRNA-resistant TBC1D4 (Fig. 6A). In a RAB10 knockdown background, insulin stimulation was unable to accelerate the mobilization of HA-GLUT4-mEos3.2 from the perinuclear region (Fig. 6B). Knockdown of RAB10 under basal conditions had no effect (Fig. 6B). Together, these data demonstrate TBC1D4 regulates mobilization of GLUT4 from the perinuclear region, and RAB10 is required for insulin-stimulated mobilization of GLUT4 from the perinuclear region.

**Figure 6.**
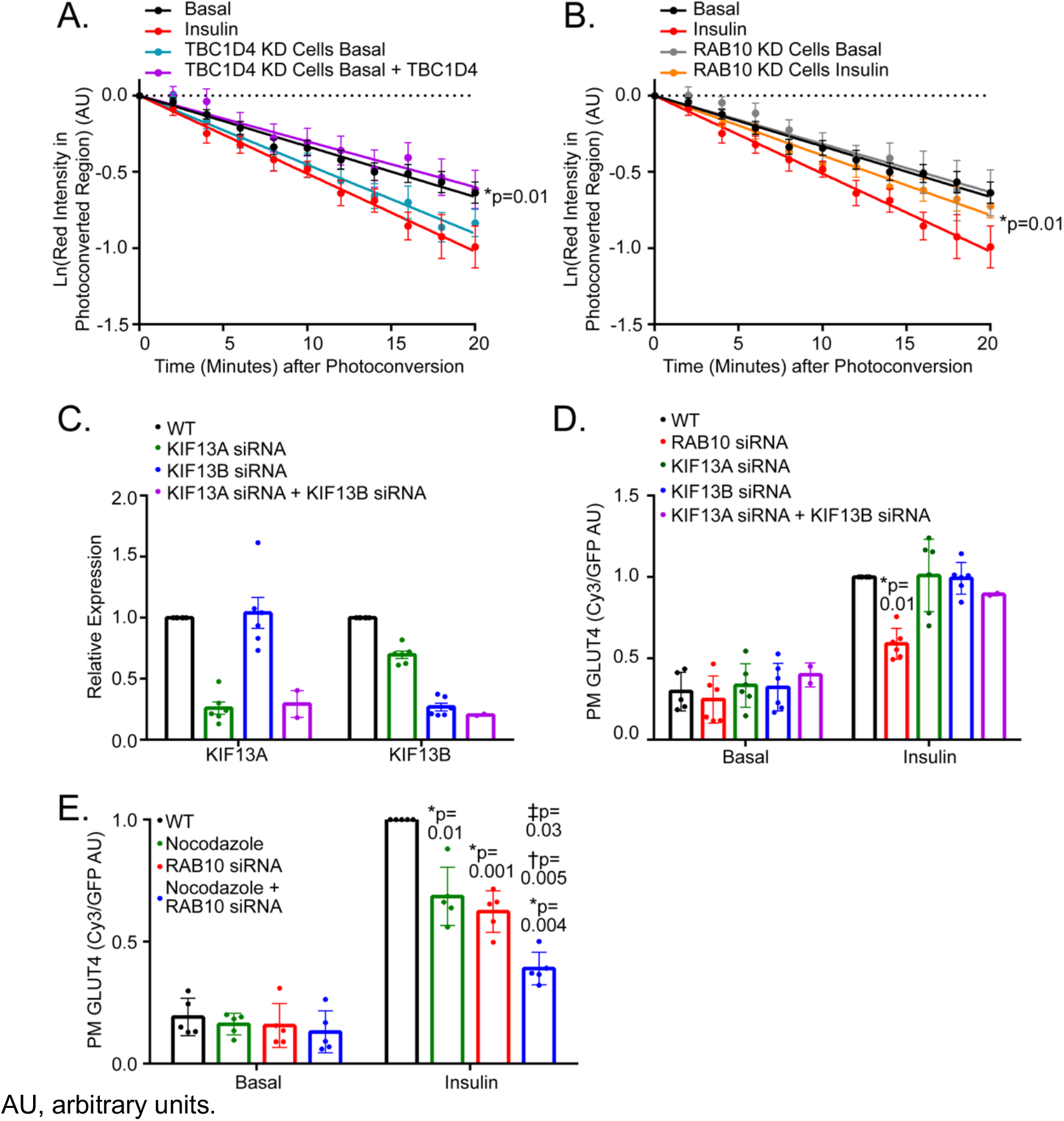
The TBC1D4-RAB10 module regulates insulin-stimulated mobilization of GLUT4 from the perinuclear region. **A.** Quantification of average red HA-GLUT4-mEos3.2 intensity in the photoconverted perinuclear region of basal live cells with stable knockdown of TBC1D4. Cells expressing exogenous TBC1D4 where indicated. Data from basal and insulin-stimulated wildtype cells (Fig. 3E) displayed. Values normalized to value at time 0. Mean normalized values ± SEM, N=3-4 assays, 5 cells per assay. *, p<0.05 comparing basal TBC1D4 KD and basal TBC1D4 KD + TBC1D4 slopes. **B.** Quantification of average red HA-GLUT4-mEos3.2 intensity in the photoconverted perinuclear region of live cells with stable knockdown of RAB10 under basal and 10nM insulin stimulated conditions. Data from basal and insulin-stimulated wildtype cells (Fig. 3E) displayed. Values normalized to value at time 0. Mean normalized values ± SEM, N=3-4 assays, 4-6 cells per assay. *, p<0.05 comparing basal RAB10 KD and insulin-stimulated RAB10 KD slopes. **C.** Quantitative RT-PCR of relative KIF13A or KIF13B mRNA expression in control 3T3-L1 adipocytes and those electroporated with KIF13A and/or KIF13B siRNAs. N=6 assays. **D.** Quantification of PM to total HA-GLUT4-GFP in serum starved cells stimulated with 1nm insulin. Values normalized to wildtype, insulin condition. N=2-6 assays ± SEM. *, p<0.05 compared to wildtype insulin-stimulated condition, two-tailed unpaired t-test, nonnormalized raw data. **E.** Quantification of PM to total HA-GLUT4-GFP in serum starved cells stimulated with 1nm insulin. siRNA targeting RAB10 electroporated where indicated, and 3μM nocodazole (or an equivalent volume of DMSO) added where indicated. Values normalized to wildtype, insulin condition. N=5 assays ± SEM. *, p<0.05 compared to wildtype, insulin condition, †, p<0.05 compared to nocodazole, insulin condition, and ‡, p<0.05 compared to RAB10 KD, insulin condition, two-tailed paired t-test, nonnormalized raw data. AU, arbitrary units.

A recent report in HeLa cells demonstrated RAB10 binding to the microtubule motor protein Kinesin 13A/B (KIF13A/B) is required for the tubulation of endosomes (28). However, in adipocytes depletion of KIF13A using 2 different siRNAs alone and in combination did not affect the amount of GLUT4 in the PM under basal or insulin-stimulated conditions compared to wildtype (WT) conditions (Fig. 6C and D). To explore the possibility that RAB10 mobilization of TGN GLUT4 in adipocytes requires a kinesin other than KIF13, we determined whether the effects of RAB10 depletion were additive to those of nocodazole-induced microtubule depolymerization on GLUT4 translocation. Nocodazole treatment resulted in a 50% decrease in the amount of GLUT4 in the PM under insulin stimulation, consistent with previous reports (2, 8, 47), and the nocodazole induced decrease in the amount of GLUT4 in the PM was additive with RAB10 (Fig. 6E). Thus, Rab10 mediated mobilization of GLUT4 from the perinuclear region does not appear to be microtubule-dependent.

## DISCUSSION

Here we show that the GLUT4 peri-nuclear storage compartment is an element of the TGN from which newly synthesized lysosomal proteins are targeted to the late endosomes and the ATP7A copper transporter is translocated to the PM by elevated copper (Fig. 7). Consequently, GLUT4 intracellular sequestration and mobilization by insulin is achieved, in part, through utilizing a region of the TGN devoted to specialized transport cargo in general rather than being specific for GLUT4. Our results define this TGN region as a sorting and storage site from which different cargo are mobilized by distinct signals.

**Figure 7.**
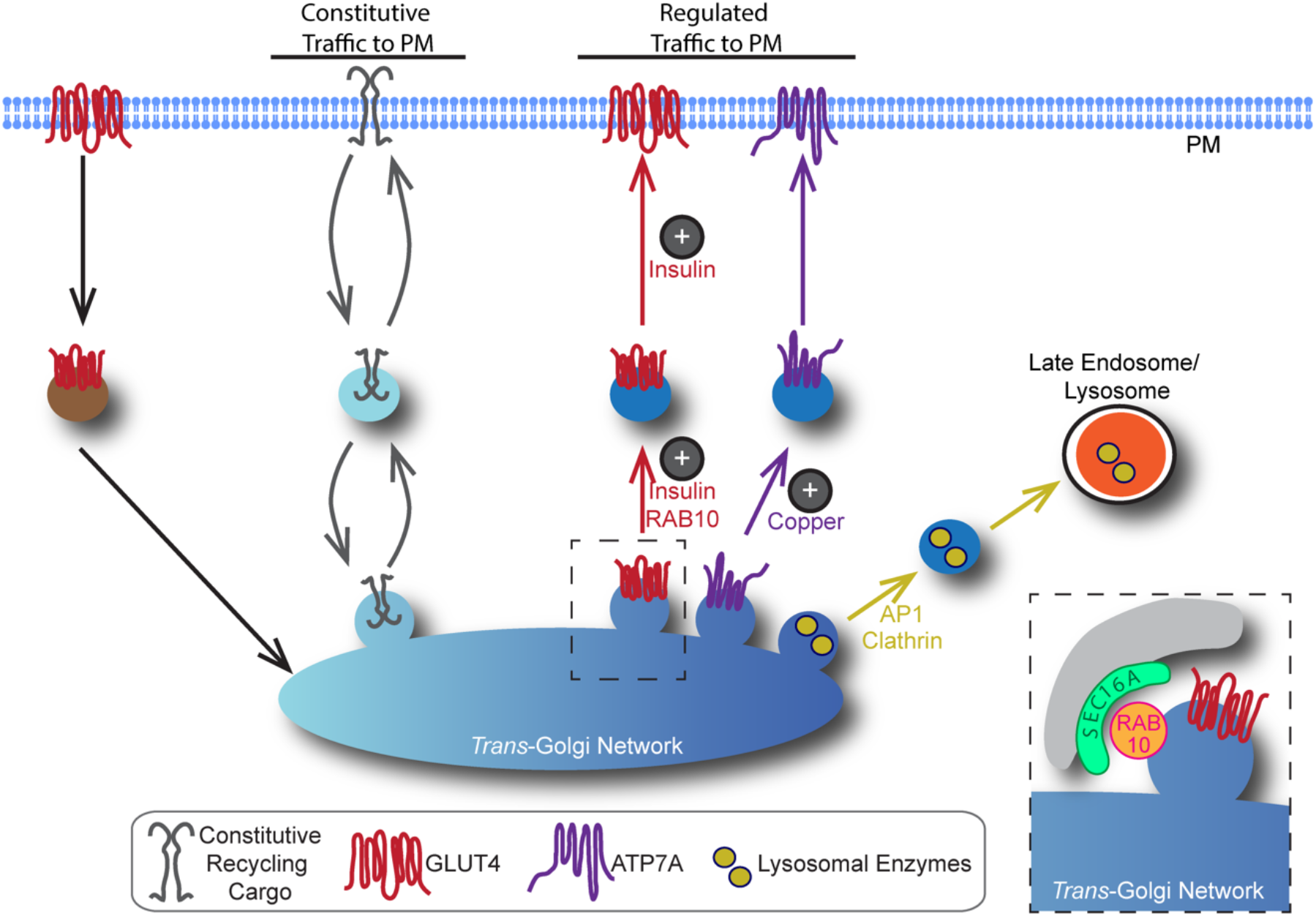
Model of GLUT4 trafficking in 3T3-L1 adipocytes. In 3T3-L1 adipocytes the biogenesis of GLUT4-containing vesicles (IRVs), copper transporter ATP7A-containing vesicles, and vesicles containing lysosomal enzymes occurs at a regulated domain of the *trans*-Golgi network (TGN); traffic of constitutive recycling proteins through the TGN occurs at an independent domain. Mobilization of ATP7A from the TGN is promoted by copper stimulation. The diversion of vesicles containing lysosomal enzymes away from traffic from the PM is mediated by the AP1 clathrin adaptor. The exocytosis of GLUT4 to the PM is accelerated by insulin. Insulin accelerates the recruitment, docking, and fusion of GLUT4-containing insulin responsive vesicles (IRVs) with the PM. Insulin also promotes the mobilization of GLUT4 from the perinuclear TGN, replenishing the IRV pool. This is important because GLUT4 in the PM is rapidly trafficked back to the TGN via the endosomal pathway. Mobilization of GLUT4 from the perinuclear region is regulated by TBC1D4, and insulin-stimulated acceleration of GLUT4 mobilization requires RAB10. Inset, SEC16A-labeled structures reside adjacent to GLUT4-containing membranes, and SEC16A organizes RAB10 at the perinuclear region.

### Insulin-stimulated acceleration of GLUT4 mobilization from the perinuclear region is regulated by the TBC1D4-RAB10 module

In this study we developed a novel photoconversion and live cell imaging assay to determine the rates of GLUT4 trafficking to and from the perinuclear compartment under different conditions. We demonstrate in intact cells that insulin accelerates the mobilization of GLUT4 from the perinuclear region by 50%. To date cell-free *in vitro* reconstitution assays using extracts of 3T3-L1 adipocytes and muscle cells have provided the best experimental evidence for insulin promoting the formation of IRVs from the TGN (10, 48). Incubation of donor membranes (i.e. the TGN) with insulin-stimulated cytosol results in an approximately 50% increase in the biogenesis of IRVs compared to incubation with basal cytosol, consistent with our results in live cells. We find insulin does not regulate GLUT4 recruitment to the perinuclear region. A large portion of GLUT4 in the PM is internalized by clathrin-mediated endocytosis together with constitutively recycling cargo such as TR (1). GLUT4-containing endosomes are sent to the TGN, and the observation that delivery of GLUT4 to the TGN is not regulated by insulin is not surprising given insulin stimulation has little effect on TR trafficking (1).

We demonstrate that the GTPase activating protein TBC1D4 and its target RAB, RAB10, are required for insulin-stimulated mobilization of GLUT4 from the perinuclear region. We find approximately 20% of BFP-RAB10 localizes to the perinuclear region and colocalizes with perinuclear GLUT4, supporting the finding that TBC1D4-RAB10 functions at the perinuclear region. Interestingly, RAB8A, the TBC1 D4-target RAB required for insulin-stimulated GLUT4 translocation in muscle, localizes to the perinuclear region in L6 muscle cells (49). We cannot exclude that RAB10 functions at the PM in addition to functioning at the TGN as suggested previously (13, 24). However, RAB10 functioning at the TGN in GLUT4 trafficking is in line with the function of RAB10 in other systems: RAB10 is involved in TLR4 trafficking from the TGN to the PM (30), membrane trafficking from the TGN required for axon development (29), and membrane transport to the primary cilia (26).

Upon insulin stimulation in 3T3-L1 adipocytes, pre-formed GLUT4 vesicles rapidly dock and fuse with the PM, increasing GLUT4 in the PM until a maximum is reached at 10 minutes of stimulation (50). GLUT4 in the PM is continually internalized and trafficked to the TGN. Accelerating the formation of IRVs that can be trafficked to the PM allows the increase in PM GLUT4 to be maintained at longer lengths of insulin stimulation (50). Expression of a dominant-negative TBC1D4 construct (TBC1D4-DN), which is mutated in four of the six Akt phosphorylation sites, blocks the RAB10 regulated GLUT4 trafficking step (20). In cells expressing TBC1D4-DN, insulin transiently increases GLUT4 in the PM within 5 minutes of stimulation, however this increase cannot be maintained at longer lengths of insulin stimulation (21). Furthermore, with TBC1D4-DN expression insulin-stimulated recruitment of GLUT4 to the PM is biphasic, with rapid exocytosis of 40% of GLUT4, followed by slow exocytosis of the remaining GLUT4 (21). The ability of insulin to initially recruit GLUT4 to the PM indicates insulin promotes the recruitment, docking, and fusion of pre-formed GLUT4 vesicles, and thus the regulation of these steps is not directly dependent on TBC1D4-RAB10. Furthermore, at basal state approximately half of GLUT4 resides in vesicles (51), consistent with the observed rapid exocytosis of 40% of GLUT4 upon insulin stimulation. The inability of insulin to maintain the initial increase in GLUT4 in the PM and the inefficient exocytosis of 50% of GLUT4 is consistent with insulin being unable to accelerate the mobilization of GLUT4 from the TGN and with TBC1D4-RAB10 regulating this step. Interestingly, in 3T3-L1 fibroblasts insulin stimulation transiently increases the amount of GLUT4 in the PM within 10 minutes of stimulation, however this increase cannot be maintained over longer lengths of stimulation (16). One explanation of these data is 3T3-L1 fibroblasts express the machinery required for insulin-stimulated increase in efficiency of IRV docking and fusion with the PM, however, they do not express the machinery required for insulin-stimulated mobilization of GLUT4 from the perinuclear compartment. The expression of such machinery may be gained throughout differentiation.

Although we have established mobilization of GLUT4 from the TGN as an insulin-controlled step dependent on TBC1D4/RAB10, we have not as yet defined the mechanism of GLUT4 mobilization. Insulin signaling could accelerate the biogenesis of IRVs at the TGN, or insulin signaling could accelerate the movement of newly formed IRVs from the perinuclear area. The latter could be accomplished by linking nascent IRVs to the cytoskeleton at the perinuclear region. The kinesin motors KIF5B (52) and KIF3 (53) have been suggested to be required for insulin-stimulated GLUT4 translocation, and RAB10 interaction with KIF13A and KIF13B has recently be shown to be required for tubulation of endosomes in HeLa cells (28). However, we find that effects of nocodazole-induced microtubule depolymerization and siRNA-mediated depletion of RAB10 on insulin-stimulated GLUT4 translocation are additive, arguing RAB10-mediated mobilization of GLUT4 from the perinuclear region is not dependent on microtubules. RAB10 has been shown to interact with the myosin motor MYO5A (54), and RAB10-MYO5A interaction has been suggested to regulate IRV docking/fusion in adipocytes (13). Interestingly, in muscle MYO5A interaction with RAB8A is argued to regulate GLUT4 trafficking at the perinuclear region (49). Furthermore, in neurons RAB10 interaction with MYO5B is required for the fission of RAB10 vesicles at the TGN (29). Thus, it may be useful to think about RAB10 possibly interacting with myosin motors to regulate IRV formation and/or link them to the cytoskeletal system. Of note, KIF13B, KIF5B, and MYO5A were present in immunoabsorbed GLUT4-containing membranes, however none were differentially immunoabsorbed in F^5^Y-GLUT4 membranes.

### SEC16A is important for RAB10 localization at the perinuclear region

Here we have advanced the understanding of the role of SEC16A in GLUT4 trafficking. We show the previously described SEC16A-labeled structures that surround GLUT4 in the perinuclear TGN (31), are also associated with RAB10. The spatial organization of perinuclear GLUT4-SEC16A-RAB10 is not random. Nocodazole depolymerization of microtubules disperses GLUT4 (45, 46), yet the organization of GLUT4-SEC16A-RAB10 is retained. The distance between adjacent peaks of GLUT4 and SEC16A is approximately 800nm with or without nocodazole treatment, and RAB10 remains colocalized with GLUT4 and SEC16A. A peak-to-peak distance of 800nm is in line with the average diameter of a Golgi cisternae, which has been calculated to range from 500-1000nm (55). We further find that siRNA-mediated depletion of SEC16A results in a 30% reduction of RAB10 in the perinuclear region under basal and insulin-stimulated states. These data argue SEC16A is important for localizing RAB10 to the perinuclear region, and SEC16A can bind to RAB10 whether it is bound to GDP or GTP. These data are consistent with the known role of SEC16A at ERES, where it acts as a scaffold for organization of COPII components (32, 33).

Mutations in the leucine-rich repeat kinase 2 (LRRK2) are associated with Parkinson’s disease (PD). LRRK2 has been suggested to regulate SEC16A localization at ERES (56). More recently LRRK2 has been shown to phosphorylate a subset of RAB proteins, including RAB10 (57). RAB10 phosphorylation status at LRRK2 sites has been implicated in regulation of ciliogenesis, and expression of mutant LRRK2 with defects in ciliogenesis (58, 59). It will be interesting to determine if PD-associated mutations in LRRK2 have any effect on GLUT4 trafficking in adipocytes.

### A role for the GLUT4-containing TGN in the biogenesis and sorting of specialized vesicular carriers

We identified the protein composition of the GLUT4-containing TGN by identifying proteins enriched in F^5^Y-GLUT4 immunoabsorption compared to WT and F^5^A-GLUT4-containing immunoabsorption. The enrichment of lysosomal enzymes known to traffic from the TGN to late endosomes/lysosomes and the ATP7A copper transporter lead us to conclude that IRVs form from a region of the TGN where unrelated transport vesicles containing other specialized cargos form. None of the specialized vesicles formed at the GLUT4-containing TGN follow the transferrin receptor (TR)-containing constitutive trafficking pathway from the TGN to the PM, suggesting GLUT4 and specialized cargo reside in a region of the TGN distinct from the region where vesicles that constitutively traffic form. We do not know if the detergent-free cell lysis method used in the immunoabsorption of GLUT4-containing compartments keeps individual stacks of the TGN intact, or if the method results in fragmentation of a TGN stack. However, the enriched immunoabsorption of cargo whose trafficking is specialized, but not constitutively recycling cargo, argues we are able to distinguish different regions or subdomains of the TGN. Interestingly, when GLUT4 is ectopically expressed in cell types that do not natively express GLUT4, such as fibroblasts, CHO cells, and HeLa cells, an insulin regulated recycling mechanism does exist, albeit less robust than in adipocytes (12, 60). Specifically, it has been demonstrated that GLUT4 travels to the PM in vesicles that are distinct from vesicles carrying constitutively recycling cargo (12). The ATP7A copper transporter is more widely expressed than is GLUT4. Hence, the specialized TGN subdomain that contains GLUT4 and ATP7A in 3T3-L1 adipocytes likely exists in other cell types that do not natively expressing GLUT4 explaining why there is rudimentary insulin-regulation of GLUT4 traffic when it is ectopically expressed in these other cell types.

Two major destinations for proteins in the TGN are the late endosome/lysosome and the PM. The mannose 6-phosphate receptor (MPR) and AP1 clathrin adaptin complex are required for diverting cargo destined for the late endosome/lysosome away from the PM (41). Their enrichment in F^5^Y-GLUT4-containing perinuclear compartments argues the GLUT4-containing TGN is the site where lysosomal enzymes are sorted into specialized transport vesicles that traffic to the late endosome/lysosome. Previous immunofluorescence and electron microscopy studies have demonstrated GLUT4 colocalizes with MPR (4) and AP1 (61), validating their enrichment. Trafficking of the copper transporter ATP7A between the TGN and the PM is known to be tightly regulated by copper load to maintain copper homeostasis (42). Enrichment of ATP7A in F^5^Y-GLUT4-containing perinuclear compartments argues the GLUT4-containing TGN is also the site where select regulated recycling membrane proteins are packaged in transport vesicles that travel to the PM. Colocalization of ATP7A with GLUT4 in the TGN is supported by the observation that with nocodazole treatment ATP7A remains colocalized with GLUT4 in a subset of fragments. ATP7A is mobilized from the GLUT4-containing TGN in response to elevated copper, but not insulin stimulation. On the other hand, copper stimulation does not induce translocation of GLUT4. These data demonstrate stimuli mobilize specific cargo from the GLUT4-containing TGN.

Proteins localized to the TGN and TGN transport vesicles were the most significantly enriched in F^5^Y-GLUT4-containing perinuclear compartments. However, there was also an enrichment of endosome, Golgi, and ER-to-Golgi intermediate compartment (ERGIC) proteins. These compartments are found in the compact perinuclear region, raising the possibility that the packaging and sorting of cargo in transport vesicles at the perinuclear region involves the interplay of membrane compartments of different natures. In human cells a clathrin heavy chain isoform, CHC22, has been proposed to function at the ERGIC to sequester newly synthesized GLUT4 in insulin responsive vesicles (IRVs) (60). Mice do not have an equivalent CHC22 gene and it has been suggested that CHC17 isoform might substitute for CHC22 in regulation of GLUT4 in mice (60). It is of interest to note that mouse clathrin heavy chain protein (CLTC)/ CHC17 was one of the most abundant proteins based on signal intensity in all 4 immunoabsorption experiments, although CLTC was not differentially absorbed in any of the comparisons. Clathrin is required for AP1-mediated vesicle trafficking between the TGN and late endosome, and therefore immunoisolation of clathrin with GLUT4 is consistent with GLUT4 localization to the region of the TGN were AP1-containing vesicles are formed.

## MATERIALS AND METHODS

### cDNA constructs, siRNA, antibodies, chemicals, and drugs

cDNA constructs encoding wild-type (WT), F^5^Y, F^5^A-HA-GLUT4-GFP, and TBC1D4 have been previously described (20, 62, 63). The HA-GLUT4-mEos3.2 cDNA construct was generated by replacing GFP in the HA-GLUT4-GFP cDNA construct for mEos3.2 (Addgene plasmid #54525) (43) through restriction cloning. KpnI and BamHI restriction sites respectively flank the N- and C-terminuses of GFP. A wobble mutation was made at an internal KpnI site in mEos3.2 to prevent its digestion using the QuikChange II XL Site-Directed Mutagenesis kit (200521; Agilent Technologies) and following primer pair: 5’-GTT CGA TTT TAT GGT ACT AAC TTT CCC GCC AAT GG-3’ and 5’-CCA TTG GCG GGA AAG TTA GTA CCA TAA AAT CGA AC-3’. mEos3.2 with the wobble internal KpnI site was PCR amplified with an N-terminal primer containing a KpnI restriction site: 5’-GCTTGGTACCATGAGTGCG-3’, and C-terminal primer containing a BamHI restriction site: 5’-GCTAGGATCCTTATCGTCTGGC-3’. The BFP-RAB10 cDNA construct was a kind gift from Gia Voeltz at University of Colorado Boulder.

Antibodies against Syntaxin6 (ab12370; Abcam and 2869T; Cell Signaling), TGN46 (ab16059; Abcam), LAMP1 (ab25630; Abcam), ATP7A (LS-C209614; LSBio), GM130 (610822; BD Transduction), SEC16A (KIAA0310; ProteinExpress), and Haemagglutinin (HA) tag (901503; Biolegend) were used for immunofluorescence.

Chemicals and drugs used were MK-2206 (11593; Cayman), Nocodazole (M1404; Sigma-Aldrich), Bathocuproinedisulfonic acid disodium salt (BCS) (B1125-500MG; Sigma-Aldrich), and Copper(II) chloride dehydrate (C3279; Sigma-Aldrich).

The siRNA constructs targeting RAB10 and SEC16A were as previously published. RAB10: si251, 5’-GCA UCA UGC UAG UGU AUGA-3’ (same sequence as shRNA expressed by RAB10 KD cells; (22)). SEC16A: si1, 5’-CTT CAG AAT ATC AGC TCC CTG GGG CTC-3’, si3, 5’-AGC TGG ACT TGC TGG TGG CTG GGC CAA-3’ (31) (two siRNA were used to target SEC16A to achieve a greater reduction in RNA). The siRNAs for KIF13A and KIF13B were designed at Integrated DNA Technologies (IDT). KIF13A: si2, 5’-ATC CTT TAA ATA GTA AAC CAG AAG CTC-3’. KIF13B: si2, 5’-CAC ATT TGG TAT GTA AGT CAA TTT CTC-3’.

### Cell lines and culture

3T3-L1 pre-adipocytes (fibroblasts) were cultured and differentiated into adipocytes as previously described (Zeigerer et al., 2002). Experiments were performed on day 5 after differentiation. 3T3-L1 adipocyte cell lines stably expressing shRNA sequences against RAB10 or TBC1D4 have been described previously (21, 22).

Immunoabsorption experiments were performed using 3T3-L1 cell lines stably expressing WT and mutant HA-GLUT4-GFP. To generate these cell lines, cDNA constructs encoding wild-type (WT), F^5^Y, and F^5^A-HA-GLUT4-GFP (62, 63) were subcloned into the pLenti6/V5-D™-TOPO^®^ vector (K4955-10; Life Technologies). 293FT packaging cells were transfected with lentiviral cDNA using Lenti-X packaging system (631276; Takara). Cultured media containing lentiviral particles was harvested after 72h and used to infect 3T3-L1 preadipocytes. HA-GLUT4-GFP-positive cells were sorted by FACS and cultured in selection medium supplemented with blasticidin (A11139-03; Invitrogen).

### Electroporation of adipocytes

Differentiated 3T3-L1 adipocytes were electroporated with 45-55μg of cDNA constructs as described previously (50). Adipocytes were electroporated with 2nmol of siRNA where indicated. When two siRNAs were used 2 nmol of each siRNA was electroporated. Assays were performed 12-72 hours post electroporation as described.

### Quantitative RT-PCR

Measurement of KIF13A and KIF13B siRNA-mediated knockdown was performed by quantitative RT-PCR. At 72 hours post electroporation, cells were harvested, RNA extracted using the RNeasy kit (74106; QIAGEN), and cDNA prepared from extracted RNA using the RNA to cDNA EcoDry Premix (639545; Takara Bio Inc.). Quantitative RT-PCR was performed using appropriate primer pairs from the PrimerBank database. Primer pair to KIF13A: Forward, 5’-TCG GAT ACG AAG GTA AAA GTT GC-3’ and Reverse, 5’-CTG CTT AGT GTT GGA AGG AGG-3’. Primer pair to KIF13B: Forward, 5’-GCT CTG TAG TGG ACT CTT TGA AC-3’ and Reverse, 5’-TTT GGG GTC AAG AAG GTC TCG-3’.

### GLUT4 translocation (surface-to-total)

HA-GLUT4-GFP has a HA-epitope engineered into the first exofacial loop and GFP fused to its cytoplasmic carboxyl domain. The amount of the reporter in the PM of individual cells was determined by anti-HA immunofluorescence, normalized to the GFP fluorescence. GLUT4 translocation assay was performed as described previously (62). Briefly, cells expressing HA-GLUT4-GFP were incubated in serum free media for 2 hours. Cells were stimulated with 1nM or 10nM insulin for 30 minutes to achieve steady state GLUT4 surface levels. Cells were fixed with 3.7% formaldehyde for 6-10 minutes, and an anti-HA antibody (901503; Biolegend) was used, without permeabilization, to label HA-GLUT4-GFP on the cell surface. HA staining was visualized with Cy3 fluorescently tagged secondary antibody (115-165-062; Jackson Immunoresearch) and total HA-GLUT4-GFP was visualized by direct fluorescence, as described later.

### Copper Transporter ATP7A mobilization assay

HA-GLUT4-GFP expressing cells were treated with 200μM Bathocuproinedisulfonic acid disodium salt (BCS) for 2 hours to achieve low copper conditions, followed by stimulation with 200μM Copper(II) chloride dehydrate for 2 hours or 1nM insulin for 30 minutes. Cells were fixed and stained for native ATP7A and Syntaxin6 in the presence of 0.5mg/ml saponin.

### HA-GLUT4-mEos3.2 Photoconversion Assay

HA-GLUT4-mEos3.2 expressing cells were serum starved for 2 hours in live cell imaging media containing Dulbecco’s Modified Eagle’s Medium (DMEM) without phenol red (D5030; Sigma-Aldrich) and supplemented with 4500 mg/L D-glucose (G7528; Sigma-Aldrich), 4mM L-glutamine (G8540; Sigma-Aldrich) 4.76 g/L HEPES (H3375; Sigma-Aldrich), 1mM sodium pyruvate (11360, Life Technologies), and 2.5g/L Sodium Bicarbonate (S6297; Sigma-Aldrich) at pH 7.2. Where indicated cells were subsequently stimulated with 10nM insulin for 10 minutes. Cells were then transferred to the confocal microscope, where they were housed in an incubation chamber at 37°C, 5% CO_2_. Set up on the scope took approximately 5 minutes once the sample was placed, thus making the total incubation time in insulin prior to photoconversion 15 minutes (at 15 minutes of insulin stimulation, cells have achieved insulin-stimulated steady state conditions). For experiments where the AKT inhibitor MK2206 was added, cells were treated with 1μM MK2206, or equivalent volume of DMSO, for the last hour of the starvation period, as well as during the 15 minute incubation period with insulin (75 minutes total).

### Microscopy, image quantification, and statistical analysis

#### Epifluorescence

Epifluorescence images were collected on an inverted microscope at room temperature using a 20x air objective (Leica Biosystems) and a cooled charge-coupled device 12-bit camera. Exposure times and image quantification (12) were performed using MetaMorph image processing software (Universal Imaging) as previously described. GFP and Cy3 fluorescence signals were background corrected and the surface(Cy3)/total(GFP) (S/T) GLUT4 was calculated for each cell. The S/T values were normalized within each assay to the mean S/T value for the indicated condition to allow for averaging results across multiple biological repeat assays. Unpaired student’s *t* tests were performed on raw (non-normalized) S/T mean values from multiple assays. To quantify the fraction of BFP-RAB10 in the perinuclear region, cells were co-transfected with HA-GLUT4-GFP. Perinuclear HA-GLUT4-GFP was used as a marker to create an outline of the perinuclear region, and the outline was transferred to the image of BFP-RAB10. As a measure of the fraction of BFP-RAB10 in the perinuclear region, the integrated BFP-RAB10 intensity in the outlined perinuclear region was calculated, and divided by the total integrated intensity of BFP-RAB10 in the cell. Unpaired student’s *t* tests were performed on raw (non-normalized) S/T mean values from multiple assays.

#### Airyscan confocal experiments

Airyscan confocal images were collected on a laser scanning microscope (LSM880; ZEISS) with Airyscan using a 63x objective. Pearson’s correlation coefficient (r) for HA-GLUT4-GFP and ATP7A was calculated with MetaMorph software by generating binary masks of HA-GLUT4-GFP and ATP7A using the 98^th^ percentile grayscale value. For quantification of native ATP7A overlap with Syntaxin6, a threshold using the 98^th^ percentile grayscale value was set on the image of native ATP7A and on the image of Syntaxin6. A binary mask of the thresholded Syntaxin6 was generated, and percent of thresholded ATP7A intensity under the mask was calculated. Unpaired student’s *t* tests were performed on individual cells from the indicated conditions.

#### Linescan analyses

Linescan plots were generated using the Linescan application in MetaMorph or Image J. Radial linescan plots were generated using the Radial Profile Plot plugin in Image J (https://imagej.nih.gov/ij/plugins/radial-profile.html). For each radial linescan plot, five HA-GLUT4-GFP fragments were selected based on high HA-GLUT4-GFP intensity. A circle with a radius of 40 pixels was applied to the fragment and centered on the peak of HA-GLUT4-GFP fluorescence. A plot of normalized integrated HA-GLUT4-GFP, SEC16A, and BFP-RAB10 fluorescence intensities around the circle (sum of integrated pixel values around circle/ total number of pixels) (y-axis) were plotted for each distance from the center of the circle (x-axis).

#### HA-GLUT4-mEos3.2 photoconversion and live cell imaging

Photoconversion of HA-GLUT4-mEos3.2 and image collection was performed on a laser scanning microscope (LSM880; ZEISS) with incubation chamber using a 63x objective. Green and red pre-photoconversion images of a cell expressing HA-GLUT4-mEos3.2 were acquired by excitation with 488nm and 561nm lasers, respectively. A high scan speed of 10, no averaging, and a low laser power of 0.2% were used to prevent photobleaching. A designated section of the perinuclear region was then bleached with a 405nm laser at 20% power, scan speed of 7 for 12 cycles. Green and red post-photoconversion images of the cell expressing HA-GLUT4-mEos3.2 were acquired every 2 minutes for a total of 20 minutes. The definite focus option was used in attempt to prevent drift in the z axis. For the average red intensity value in the photoconverted region at each time point post-photoconversion, the pre-photoconversion red intensity value was subtracted in MetaMorph. Values were then normalized to the 0 minute post-photoconversion value and the natural log taken. After averaging across multiple cells, a linear curve fit was applied. Statistical comparison of slopes was performed in Prism by calculating a two-tailed p value from testing the null hypothesis that the slopes are identical. For the average green intensity value at each time point post-photoconversion, the pre-photoconversion green intensity value was subtracted. Values were then normalized to the negative value of the 0 minute time point post-photoconversion, and added to 1. After averaging across multiple cells, an exponential curve fit was applied.

#### Immuno-isolation of native GLUT4-containing compartments, SILAC mass spectrometry, and data processing

Each pair-wise comparison was performed in inverted Forward and Reverse labeling conditions.

For the WTvsF^5^Y comparison, the labeling design was i) Forward condition: WT HA-GLUT4-GFP cells grown in light SILAC medium versus F^5^Y-HA-GLUT4-GFP cells grown in heavy SILAC medium and ii) Reverse condition: WT HA-GLUT4-GFP cells grown in heavy SILAC medium versus F^5^Y-HA-GLUT4-GFP cells grown in light SILAC medium.

For the F^5^YvsF^5^A comparison, the labeling design was i) Forward condition: F^5^Y-HA-GLUT4-GFP cells grown in light SILAC medium versus F^5^A-HA-GLUT4-GFP cells grown in heavy SILAC medium and ii) Reverse condition: F^5^Y-HA-GLUT4-GFP cells grown in heavy SILAC medium versus F^5^A-HA-GLUT4-GFP cells grown in light SILAC medium.

The objective of this experiment was to identify by SILAC mass spectrometry proteins colocalized with GLUT4 in the perinuclear compartment based on enrichment with F^5^Y-GLUT4 in immunoabsorption, not to identify all proteins in GLUT4-containing compartments. Therefore, we did not include a control condition to identify proteins that are non-specifically absorbed during the immunoisolation.

#### Stable isotope labeling of cultured cells

Stable HA-GLUT4-GFP-expressing 3T3-L1 preadipocytes were grown for 5 doublings and differentiated in Lysine (LYS) and Arginine (ARG)-deficient DMEM (89985; Thermo Scientific), supplemented with 10% dialyzed FBS, and 42ug/ml of either LYS-HCL and ARG-HCL normal isotopes (Light SILAC medium) or with ^13^C_6_LYS and ^13^C_6_LYS,^15^N4 ARG isotopes (Heavy SILAC medium) (89983 and 88210; Pierce) at 37°C in 5% CO_2_. Under these conditions, the isotopes incorporation efficiency was higher than 95%, without detectable arginine to proline conversion.

#### Immuno-isolation of GLUT4-containing compartments

Day 5 post-differentiation, labeled stable HA-GLUT4-GFP-expressing 3T3-L1 adipocytes were incubated in serum-free either Light or Heavy SILAC media for 2 h at 37°C in 5% CO_2_ to establish basal GLUT4 retention. Cells were washed one time with PBS, harvested into 1 ml of HES buffer (20mM HEPES, 1mM EDTA, 250mM sucrose, and protease inhibitors) and homogenized by subsequent passage through 22G^1/2^ and 27G^1/2^ syringes on ice. Total cell homogenates were cleared by successive centrifugations at 1000g for 10 minutes to remove unbroken cells, nuclei and fat. Protein concentration of both Light-cultured cells and Heavy-cultured cells was measured by BCA assay (23225; Thermo Scientific) and homogenates were mixed to a 1:1 ratio. HA-GLUT4-GFP-containing compartments were isolated by incubation for 30 minutes at 4°C with magnetic GFP-bound beads (130-091-125; Miltenyi Biotech). Beads were washed 5 times in PBS supplemented with protease inhibitors and absorbed material was eluted with elution buffer (50mM Tris HCl (pH6.08), 50 mM DTT, 1%SDS, 1nM EDTA, 0.005% bromophenol blue, 10% glycerol).

#### LC-MS/MS and Bioinformatics analysis

Eluates were resolved on 5-20% gradient SDS-Page gel and subjected to in-gel digest followed by LC-MS/MS analysis as described (64). Peptide/spectrum matching as well as false discovery control (1% on the peptide and protein levels, both) and protein quantitation were performed using the MaxQuant suite of algorithms (65). We used the SILAC ratio of polypeptides in the immunoabsorbates to identify proteins enriched with F^5^Y-GLUT4. The comparisons of F^5^Y to WT and F^5^Y to F^5^A were performed twice, switching which sample was labeled with heavy amino acids. We identified the proteins whose average ratios in the two F^5^Y vs WT and F^5^Y vs F^5^A experiments were greater than 1.3 fold with a ‘significance B’ (65) < 0.05, falling back on the method due to the small n. There were 508 proteins enriched in the F^5^Y vs the combined WT and F^5^A data sets. We used the merged data set for downstream computational analyses.

## AUTHOR CONTRIBUTIONS

A. Brumfield, N. Chaudhary, D. Molle, and J. Wen designed, performed, and analyzed experiments. J. Graumann performed mass spectrometry. A. Brumfield and T.E. McGraw wrote the manuscript. T.E. McGraw conceived of the project, designed and analyzed experiments, and supervised the project.

## ACKNOWLEDGEMENTS

We thank Leona Cohen-Gould and Sushmita Mukherjee at the Weill Cornell Medicine Optical Microscopy Core where confocal microscopy was performed, and Harold Skip Ralph at the Weill Cornell Medicine Automated Optical Microscopy Core for help with image analysis. We thank Gus Lienhard (Dartmouth Medical School), Maria Belen Picatoste Botija, Rosemary Leahey, Anudari Letian, Anuttoma Ray, Lucie Yammine, and Eyoel Yemanaberhan for helpful discussions and critically reading the manuscript. This research was supported by NIH RO1 DK52852 to T.E.M, NIH 5T32 GM008539 (AB), and an American Diabetes Association mentor-based fellowship award to TEM.

## ABBREVIATIONS LIST

GLUT4: Glucose Transporter 4,
ATP7A: Menkes Copper-Transporting ATPase,
PM: Plasma Membrane,
TGN: *Trans*-Golgi Network,
ERC: Endocytic Recycling Compartment,
ERGIC: ER-to-Golgi Intermediate Compartment,
ERES: Endoplasmic Reticulum Exit Site,
IRVs: Insulin-responsive vesicles,
TR: Transferrin receptor,
IF: Immunofluorescence,
STX6: Syntaxin6,
TGN46: *Trans*-Golgi Network Protein 2,

## REFERENCES

1. Klip A, McGraw TE, James DE. Thirty sweet years of GLUT4. The Journal of biological chemistry. 2019;294(30):11369–81.

2. Karylowski O, Zeigerer A, Cohen A, McGraw TE. GLUT4 is retained by an intracellular cycle of vesicle formation and fusion with endosomes. Molecular biology of the cell. 2004;15(2):870–82.

3. Martin OJ, Lee A, McGraw TE. GLUT4 distribution between the plasma membrane and the intracellular compartments is maintained by an insulin-modulated bipartite dynamic mechanism. The Journal of biological chemistry. 2006;281(1):484–90.

4. Martin S, Millar CA, Lyttle CT, Meerloo T, Marsh BJ, Gould GW, et al. Effects of insulin on intracellular GLUT4 vesicles in adipocytes: evidence for a secretory mode of regulation. Journal of cell science. 2000;113 Pt 19:3427–38.

5. Foster LJ, Li D, Randhawa VK, Klip A. Insulin accelerates inter-endosomal GLUT4 traffic via phosphatidylinositol 3-kinase and protein kinase B. The Journal of biological chemistry. 2001;276(47):44212–21.

6. Shewan AM, van Dam EM, Martin S, Luen TB, Hong W, Bryant NJ, et al. GLUT4 recycles via a trans-Golgi network (TGN) subdomain enriched in Syntaxins 6 and 16 but not TGN38: involvement of an acidic targeting motif. Molecular biology of the cell. 2003;14(3):973–86.

7. Li Lin V, Bakirtzi K, Watson Robert T, Pessin Jeffrey E, Kandror Konstantin V. The C-terminus of GLUT4 targets the transporter to the perinuclear compartment but not to the insulin-responsive vesicles. Biochemical Journal. 2009;419(1):105–13.

8. Foley KP, Klip A. Dynamic GLUT4 sorting through a syntaxin-6 compartment in muscle cells is derailed by insulin resistance-causing ceramide. Biology Open. 2014;3(5):314–25.

9. Jedrychowski MP, Gartner CA, Gygi SP, Zhou L, Herz J, Kandror KV, et al. Proteomic analysis of GLUT4 storage vesicles reveals LRP1 to be an important vesicle component and target of insulin signaling. The Journal of biological chemistry. 2010;285(1):104–14.

10. Xu Z, Kandror KV. Translocation of small preformed vesicles is responsible for the insulin activation of glucose transport in adipose cells. Evidence from the in vitro reconstitution assay. The Journal of biological chemistry. 2002;277(50):47972–5.

11. Larance M, Ramm G, Stockli J, van Dam EM, Winata S, Wasinger V, et al. Characterization of the role of the Rab GTPase-activating protein AS160 in insulin-regulated GLUT4 trafficking. The Journal of biological chemistry. 2005;280(45):37803–13.

12. Lampson MA, Schmoranzer J, Zeigerer A, Simon SM, McGraw TE. Insulin-regulated release from the endosomal recycling compartment is regulated by budding of specialized vesicles. Molecular biology of the cell. 2001;12(11):3489–501.

13. Chen Y, Wang Y, Zhang J, Deng Y, Jiang L, Song E, et al. Rab10 and myosin-Va mediate insulin-stimulated GLUT4 storage vesicle translocation in adipocytes. The Journal of cell biology. 2012;198(4):545–60.

14. Blot V, McGraw TE. Molecular mechanisms controlling GLUT4 intracellular retention. Molecular biology of the cell. 2008;19(8):3477–87.

15. Piper RC, Tai C, Kulesza P, Pang S, Warnock D, Baenziger J, et al. GLUT-4 NH2 terminus contains a phenylalanine-based targeting motif that regulates intracellular sequestration. The Journal of cell biology. 1993;121(6):1221–32.

16. Govers R, Coster AC, James DE. Insulin increases cell surface GLUT4 levels by dose dependently discharging GLUT4 into a cell surface recycling pathway. Mol Cell Biol. 2004;24(14):6456–66.

17. Gonzalez E, McGraw TE. Insulin signaling diverges into Akt-dependent and - independent signals to regulate the recruitment/docking and the fusion of GLUT4 vesicles to the plasma membrane. Molecular biology of the cell. 2006;17(10):4484–93.

18. Xiong W, Jordens I, Gonzalez E, McGraw TE. GLUT4 is sorted to vesicles whose accumulation beneath and insertion into the plasma membrane are differentially regulated by insulin and selectively affected by insulin resistance. Molecular biology of the cell. 2010;21(8):1375–86.

19. Bai L, Wang Y, Fan J, Chen Y, Ji W, Qu A, et al. Dissecting multiple steps of GLUT4 trafficking and identifying the sites of insulin action. Cell metabolism. 2007;5(1):47–57.

20. Sano H, Kane S, Sano E, Miinea CP, Asara JM, Lane WS, et al. Insulin-stimulated phosphorylation of a Rab GTPase-activating protein regulates GLUT4 translocation. The Journal of biological chemistry. 2003;278(17):14599–602.

21. Eguez L, Lee A, Chavez JA, Miinea CP, Kane S, Lienhard GE, et al. Full intracellular retention of GLUT4 requires AS160 Rab GTPase activating protein. Cell metabolism. 2005;2(4):263–72.

22. Sano H, Eguez L, Teruel MN, Fukuda M, Chuang TD, Chavez JA, et al. Rab10, a target of the AS160 Rab GAP, is required for insulin-stimulated translocation of GLUT4 to the adipocyte plasma membrane. Cell metabolism. 2007;5(4):293–303.

23. Lansey MN, Walker NN, Hargett SR, Stevens JR, Keller SR. Deletion of Rab GAP AS160 modifies glucose uptake and GLUT4 translocation in primary skeletal muscles and adipocytes and impairs glucose homeostasis. American journal of physiology Endocrinology and metabolism. 2012;303(10):E1273–86.

24. Sadacca LA, Bruno J, Wen J, Xiong W, McGraw TE. Specialized sorting of GLUT4 and its recruitment to the cell surface are independently regulated by distinct Rabs. Molecular biology of the cell. 2013;24(16):2544–57.

25. Vazirani RP, Verma A, Sadacca LA, Buckman MS, Picatoste B, Beg M, et al. Disruption of Adipose Rab10-Dependent Insulin Signaling Causes Hepatic Insulin Resistance. Diabetes. 2016;65(6):1577–89.

26. Babbey CM, Bacallao RL, Dunn KW. Rab10 associates with primary cilia and the exocyst complex in renal epithelial cells. American journal of physiology Renal physiology. 2010;299(3):F495–506.

27. Isabella AJ, Horne-Badovinac S. Rab10-Mediated Secretion Synergizes with Tissue Movement to Build a Polarized Basement Membrane Architecture for Organ Morphogenesis. Developmental cell. 2016;38(1):47–60.

28. Etoh K, Fukuda M. Rab10 regulates tubular endosome formation through KIF13A and KIF13B motors. Journal of cell science. 2019;132(5).

29. Liu Y, Xu XH, Chen Q, Wang T, Deng CY, Song BL, et al. Myosin Vb controls biogenesis of post-Golgi Rab10 carriers during axon development. Nature communications. 2013;4:2005.

30. Wang D, Lou J, Ouyang C, Chen W, Liu Y, Liu X, et al. Ras-related protein Rab10 facilitates TLR4 signaling by promoting replenishment of TLR4 onto the plasma membrane. Proceedings of the National Academy of Sciences of the United States of America. 2010;107(31):13806–11.

31. Bruno J, Brumfield A, Chaudhary N, Iaea D, McGraw TE. SEC16A is a RAB10 effector required for insulin-stimulated GLUT4 trafficking in adipocytes. The Journal of cell biology. 2016;214(1):61–76.

32. Sprangers J, Rabouille C. SEC16 in COPII coat dynamics at ER exit sites. Biochemical Society transactions. 2015;43(1):97–103.

33. Whittle JR, Schwartz TU. Structure of the Sec13-Sec16 edge element, a template for assembly of the COPII vesicle coat. The Journal of cell biology. 2010;190(3):347–61.

34. Blot V, McGraw TE. Use of quantitative immunofluorescence microscopy to study intracellular trafficking: studies of the GLUT4 glucose transporter. Methods Mol Biol. 2008;457:347–66.

35. Ong S-E, Blagoev B, Kratchmarova I, Kristensen DB, Steen H, Pandey A, et al. Stable Isotope Labeling by Amino Acids in Cell Culture, SILAC, as a Simple and Accurate Approach to Expression Proteomics. Molecular & Cellular Proteomics. 2002;1(5):376–86.

36. Ibarrola N, Kalume DE, Gronborg M, Iwahori A, Pandey A. A proteomic approach for quantitation of phosphorylation using stable isotope labeling in cell culture. Anal Chem. 2003;75(22):6043–9.

37. Garza LA, Birnbaum MJ. Insulin-responsive aminopeptidase trafficking in 3T3-L1 adipocytes. The Journal of biological chemistry. 2000;275(4):2560–7.

38. Morris NJ, Ross SA, Lane WS, Moestrup SK, Petersen CM, Keller SR, et al. Sortilin is the major 110-kDa protein in GLUT4 vesicles from adipocytes. The Journal of biological chemistry. 1998;273(6):3582–7.

39. Consortium TGO. The Gene Ontology Resource: 20 years and still GOing strong. Nucleic acids research. 2019;47(D1):D330–d8.

40. Ashburner M, Ball CA, Blake JA, Botstein D, Butler H, Cherry JM, et al. Gene ontology: tool for the unification of biology. The Gene Ontology Consortium. Nature genetics. 2000;25(1):25–9.

41. Braulke T, Bonifacino JS. Sorting of lysosomal proteins. Biochim Biophys Acta. 2009;1793(4):605–14.

42. Petris MJ, Mercer JF, Culvenor JG, Lockhart P, Gleeson PA, Camakaris J. Ligand-regulated transport of the Menkes copper P-type ATPase efflux pump from the Golgi apparatus to the plasma membrane: a novel mechanism of regulated trafficking. The EMBO journal. 1996;15(22):6084–95.

43. Zhang M, Chang H, Zhang Y, Yu J, Wu L, Ji W, et al. Rational design of true monomeric and bright photoactivatable fluorescent proteins. Nature methods. 2012;9(7):727–9.

44. Tan S, Ng Y, James DE. Next-generation Akt inhibitors provide greater specificity: effects on glucose metabolism in adipocytes. The Biochemical journal. 2011;435(2):539–44.

45. Cole NB, Sciaky N, Marotta A, Song J, Lippincott-Schwartz J. Golgi dispersal during microtubule disruption: regeneration of Golgi stacks at peripheral endoplasmic reticulum exit sites. Molecular biology of the cell. 1996;7(4):631–50.

46. Thyberg J, Moskalewski S. Role of microtubules in the organization of the Golgi complex. Exp Cell Res. 1999;246(2):263–79.

47. Olson AL, Trumbly AR, Gibson GV. Insulin-mediated GLUT4 translocation is dependent on the microtubule network. The Journal of biological chemistry. 2001;276(14):10706–14.

48. Kristiansen S, Richter EA. GLUT4-containing vesicles are released from membranes by phospholipase D cleavage of a GPI anchor. American journal of physiology Endocrinology and metabolism. 2002;283(2):E374–82.

49. Sun Y, Chiu TT, Foley KP, Bilan PJ, Klip A. Myosin Va mediates Rab8A-regulated GLUT4 vesicle exocytosis in insulin-stimulated muscle cells. Molecular biology of the cell. 2014;25(7):1159–70.

50. Zeigerer A, Lampson MA, Karylowski O, Sabatini DD, Adesnik M, Ren M, et al. GLUT4 retention in adipocytes requires two intracellular insulin-regulated transport steps. Molecular biology of the cell. 2002;13(7):2421–35.

51. Roccisana J, Sadler JB, Bryant NJ, Gould GW. Sorting of GLUT4 into its insulin-sensitive store requires the Sec1/Munc18 protein mVps45. Molecular biology of the cell. 2013;24(15):2389–97.

52. Semiz S, Park JG, Nicoloro SM, Furcinitti P, Zhang C, Chawla A, et al. Conventional kinesin KIF5B mediates insulin-stimulated GLUT4 movements on microtubules. The EMBO journal. 2003;22(10):2387–99.

53. Imamura T, Huang J, Usui I, Satoh H, Bever J, Olefsky JM. Insulin-induced GLUT4 translocation involves protein kinase C-lambda-mediated functional coupling between Rab4 and the motor protein kinesin. Mol Cell Biol. 2003;23(14):4892–900.

54. Roland JT, Bryant DM, Datta A, Itzen A, Mostov KE, Goldenring JR. Rab GTPase-Myo5B complexes control membrane recycling and epithelial polarization. Proceedings of the National Academy of Sciences of the United States of America. 2011;108(7):2789–94.

55. Klumperman J. Architecture of the mammalian Golgi. Cold Spring Harb Perspect Biol. 2011;3(7).

56. Cho HJ, Yu J, Xie C, Rudrabhatla P, Chen X, Wu J, et al. Leucine-rich repeat kinase 2 regulates Sec16A at ER exit sites to allow ER-Golgi export. The EMBO journal. 2014;33(20):2314–31.

57. Steger M, Tonelli F, Ito G, Davies P, Trost M, Vetter M, et al. Phosphoproteomics reveals that Parkinson’s disease kinase LRRK2 regulates a subset of Rab GTPases. Elife. 2016;5.

58. Lara Ordonez AJ, Fernandez B, Fdez E, Romo-Lozano M, Madero-Perez J, Lobbestael E, et al. RAB8, RAB10 and RILPL1 contribute to both LRRK2 kinase-mediated centrosomal cohesion and ciliogenesis deficits. Hum Mol Genet. 2019;28(21):3552–68.

59. Dhekne HS, Yanatori I, Gomez RC, Tonelli F, Diez F, Schule B, et al. A pathway for Parkinson’s Disease LRRK2 kinase to block primary cilia and Sonic hedgehog signaling in the brain. Elife. 2018;7.

60. Camus SM, Camus MD, Figueras-Novoa C, Boncompain G, Sadacca LA, Esk C, et al. CHC22 clathrin mediates traffic from early secretory compartments for human GLUT4 pathway biogenesis. The Journal of cell biology. 2020;219(1).

61. Martin S, Ramm G, Lyttle CT, Meerloo T, Stoorvogel W, James DE. Biogenesis of insulin-responsive GLUT4 vesicles is independent of brefeldin A-sensitive trafficking. Traffic. 2000;1(8):652–60.

62. Lampson MA, Racz A, Cushman SW, McGraw TE. Demonstration of insulin-responsive trafficking of GLUT4 and vpTR in fibroblasts. Journal of cell science. 2000;113 (Pt 22):4065–76.

63. Blot V, McGraw TE. GLUT4 is internalized by a cholesterol-dependent nystatin-sensitive mechanism inhibited by insulin. The EMBO journal. 2006;25(24):5648–58.

64. Graumann J, Hubner NC, Kim JB, Ko K, Moser M, Kumar C, et al. Stable isotope labeling by amino acids in cell culture (SILAC) and proteome quantitation of mouse embryonic stem cells to a depth of 5,111 proteins. Mol Cell Proteomics. 2008;7(4):672–83.

65. Cox J, Mann M. MaxQuant enables high peptide identification rates, individualized p.p.b.-range mass accuracies and proteome-wide protein quantification. Nat Biotechnol. 2008;26(12):1367–72.

